# Leveraging zebrafish embryo phenotypic observations to advance data-driven analyses in toxicology

**DOI:** 10.1101/2024.10.11.617795

**Authors:** Paul Michaelis, Nils Klüver, Silke Aulhorn, Hannes Bohring, Jan Bumberger, Kristina Haase, Tobias Kuhnert, Eberhard Küster, Janet Krüger, Till Luckenbach, Riccardo Massei, Lukas Nerlich, Sven Petruschke, Thomas Schnicke, Anton Schnurpel, Stefan Scholz, Nicole Schweiger, Daniel Sielaff, Wibke Busch

## Abstract

Zebrafish have emerged as a central model organism in toxicological research. Zebrafish embryos are exempt from certain animal testing regulations which facilitates their use in toxicological testing. Next to the zebrafish embryo acute toxicity test (ZFET) according to the OECD TG 236, fish embryos are used in mechanistic investigations, chemical screenings, in ecotoxicology, and drug development. However, inconsistencies in the applied test protocols and the monitored endpoints in addition to a lack of standardized data formats, impede comprehensive meta-analyses and cross-study comparisons. To address these challenges, we developed the Integrated Effect Database for Toxicological Observations (INTOB), a comprehensive data management tool that standardizes collection of metadata and phenotypic observations using a controlled vocabulary. By incorporating data from more than 600 experiments into the database and subsequent comprehensive data analyses, we demonstrate its utility in improving the comparability and interoperability of toxicity data. Our results show that the ZFET can detect toxicity spanning seven orders of magnitude at the scale of effect concentrations. We also highlight the potential of read-across analyses based on morphological fingerprints and their connection to chemical modes of action, provide information on control variability of the ZFET, and highlight the importance of time for mechanistic understanding in chemical exposure-effect assessments. We provide the full FAIR dataset as well as the analysis workflow and demonstrate how professional data management, as enabled with INTOB, marks a significant advancement by offering a comprehensive framework for the systematic use of zebrafish embryo toxicity data, thus paving the way for more reliable, data-driven chemical risk assessment.

**Synopsis:** This article shows how a novel data management tool for zebrafish embryo toxicity data advances comparative chemical risk assessment by enhancing data and metadata standardization, comparability, and cross-study analyses.

## Introduction

The release of chemicals into the environment due to human activities poses significant risks to both environmental and human health. Consequently, different legislations worldwide request comprehensive risk assessment data for the registration of industrial chemicals, pesticides, biocides, and pharmaceuticals. This includes directives and guidelines from various regulatory bodies (EPA, 1976; EU, 2009, 2008, 2006). The required data encompass toxicity information across different trophic levels and necessitate experimental testing with vertebrates, particularly fish. Several methods are particularly effective for determining toxicity thresholds and concentrations, such as estimating lethal concentration (LCx) and effective concentration (ECx) values or deriving points of departure (POD). As there is great societal and ethical demand to replace animal testing the framework of the 3Rs (Replacement, Reduction, Refinement) for animal testing was established. Furthermore, there has been a shift towards information-driven evidence-based risk assessment for humans and the environment using more mechanistic information, e.g. within the framework of the adverse outcome pathways (AOP, (Ankley et al., 2010; Patlewicz et al., 2013)).

Zebrafish (*Danio rerio*) are increasingly recognized as a valuable model for studying chemical-induced toxicity, not only in environmental toxicology but also in human health. As zebrafish possess orthologs for 70% of human genes, 80% of human disease-related genes, and 86% of general human drug targets (Gunnarsson et al., 2008; Howe et al., 2013), the model represents a powerful translational system for human hazard and risk assessment, and disease models in drug discovery (Knudsen et al., 2013; Patton et al., 2021; Peterson and Macrae, 2012; Tal et al., 2020; Truong et al., 2014). Zebrafish are conveniently maintained and bred in laboratories, and their development is rapid and observable due to the translucency of the egg chorion. They undergo external fertilization and have high reproduction rates, allowing for waterborne exposure to compounds and enabling medium to high throughput experiments with embryos. Additionally, the small sized and transparent embryos permit direct observation of developmental delays and malformations. Furthermore, up to 5 days post-fertilization zebrafish embryos (ZFE) are considered as non-protected stages and as an alternative to animal testing under European legislation (EU Directive 63, 2010; Strähle et al., 2012). This offers the development of new approach methods (NAMs) using the zebrafish embryo as whole organism model in certain regulatory frameworks (De Castelbajac et al., 2023; Tal et al., 2024).

The zebrafish embryo acute toxicity test (ZFET) has already been standardized (OECD, 2013) and has been promoted as an alternative method for the acute fish toxicity test (AFT). The ZFET can be used in weight-of-evidence approaches within Registration, Evaluation, Authorization, and Restriction (REACH) regulations and it is already used as an approved method for routine testing of waste water effluents (Scholz et al., 2013, 2008). The ZFE also served as model system in developmental toxicity studies to detect teratogenicity and/or developmental neurotoxicity of chemicals (d’Amora and Giordani, 2018; McCollum et al., 2011). Furthermore, detailed responses on different effect levels, e.g. morphological phenotype, behavior, or abundance of genes and proteins at the molecular level determined after chemical exposures of ZFE have been used to identify and define toxicological effect fingerprints for particular modes of chemical action (Kokel et al., 2010; Schüttler et al., 2019; Teixidó et al., 2022; Truong et al., 2020, 2014).

The use of the ZFE model within these different research areas and its use within the regulatory context, have increased the number of scientific publications and respective observational data. However, data often do not comply with the Findable, Accessible, Interoperable and Reusable (FAIR) principles. For example, they are often provided as analyzed results for a particular research hypothesis but not as raw data. Furthermore, varying nomenclature is used in scientific published literature for toxicological effects of chemicals using early-life stages of zebrafish. These inconsistencies underscore the need for harmonizing terminology, experimental metadata, and observational data. It has been shown that providing phenotypic ontology terms and harmonizing data and descriptions led to an improvement of results and the interoperability of data (Thessen et al., 2022). Partially, information of chemical effects on zebrafish have been added to manually curated, publicly available databases like the ECOTOX database (https://cfpub.epa.gov/ecotox/) and the zebrafish information network (ZFIN, https://zfin.org/). However, missing detailed information and metadata, e.g. on exposure conditions and the lack of standardized ontologies prevent data from being analyzed jointly, compared with each other, or used in interoperable ways, e.g. for model training (Bradford et al., 2022; Olker et al., 2022).

To overcome the above-mentioned obstacles in comparability and joint assessment of toxicological phenotypes we developed an integrated effect data base for toxicological observations (INTOB) for ZFE toxicity data. The tool supports the documentation of experimental metadata and phenotypic observations with a defined vocabulary for observed toxicological endpoints in a structured and machine-readable format, facilitating joint analyses incorporating data from many experiments. Next to applying the software in performing the ZFET in our laboratories since 2020, we also entered ZFET data of the last 20 years into the database that were available and stored at our institute. Based on the available data, which we provide via Zenodo, we performed analyses that provide insights into ZFE control variability, correlations of endpoints over time, and similarities of effect fingerprints across chemicals. A comparison with data from literature reveals that data retrieved from literature are, so far, unsuitable to perform meta-analyses of ZFET data, underlining the value of the established INTOB system.

## Methods

### Overall Approach

Our modular data management tool INTOB provides a relational database structure and a web-based user interface for data entry and enables to follow the FAIR (Findable, Accessible, Interoperable and Reusable) principles for zebrafish embryo (ZFE) toxicity data. The system allows a structured storage of phenotypic observations at different time points during the exposure experiments, and facilitates data export for subsequent analysis (Figure 1). The experimental metadata entries were defined and entered according to Supplementary Table 1. Next to an upload option for observations from automated image analysis, a defined vocabulary was used for microscopic observations with ZFE at different stages (Table 1) including the terms “coagulation”, “lack of somite formation”, “non-detachment of the tail”, “lack of heartbeat” of the OECD TG 236 (OECD, 2013). We recorded observations for 638 experiments and analyzed them with regard to data quality, effect concentrations and phenotypic effect patterns.

**Figure 1:**
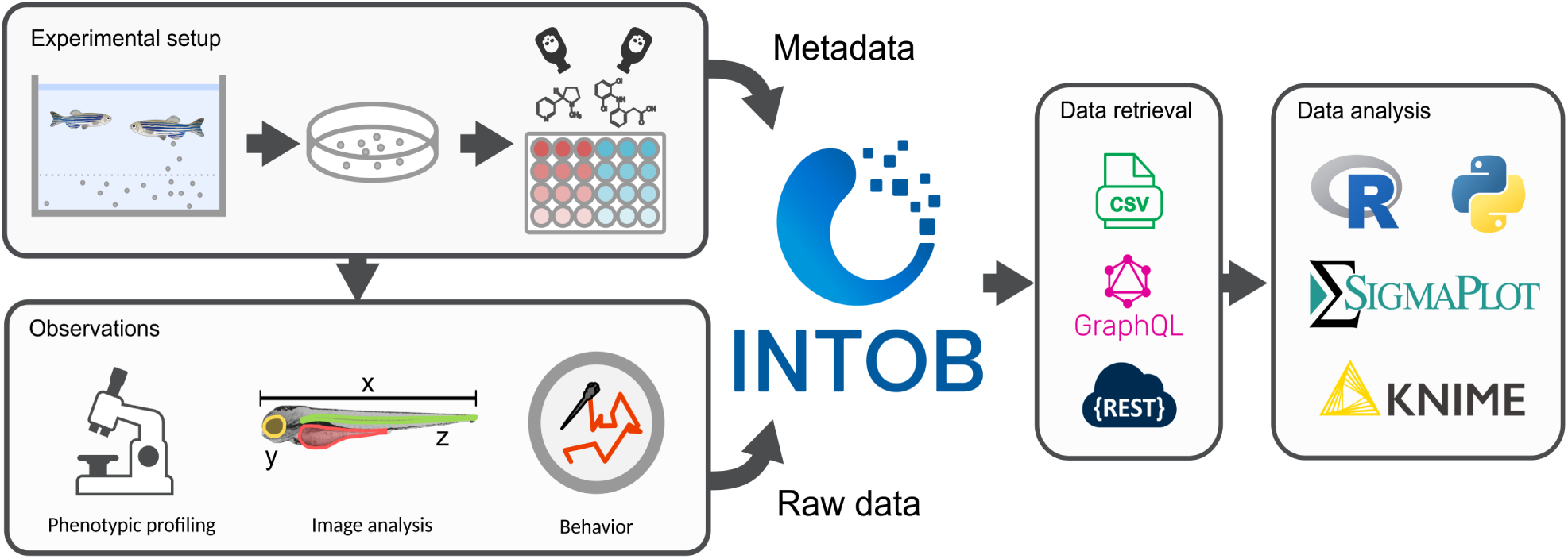
Data storage and retrieval structure of INTOB. Experimental metadata is stored alongside observations. Data can be retrieved via the user interface in csv format or an application programming interface (API).

**Table 1:** List of morphological effects with unified vocabulary. The time window where this effect can be observed is defined. Effects are organized in a tree structure, where a parent effect can entail several more detailed related effects. The effect names „coagulated“, „lack of somites“, „no tail detachment“ and „no heartbeat“ are compliant with those specified in OECD TG 236.

| effect id | parent id | Effect | Time point start (hpf) | Time point end (hpf) |
| --- | --- | --- | --- | --- |
| 1 | - | no embryo | 0 | 120 |
| 2 | - | coagulated | 0 | 120 |
| 3 | - | normal | 0 | 120 |
| 4 | - | abnormal eye | 0 | 120 |
| 5 | - | abnormal tail effects | 0 | 120 |
| 6 | - | developmental delay effects | 0 | 120 |
| 7 | - | deformation / malformation | 0 | 120 |
| 8 | - | abnormal behavior | 0 | 120 |
| 9 | - | heartbeat | 48 | 120 |
| 10 | - | abnormal blood circulation | 48 | 120 |
| 11 | - | edema | 24 | 120 |
| 12 | - | abnormal pigmentation | 48 | 120 |
| 13 | - | abnormal hatching | 48 | 120 |
| 14 | - | abnormal swim bladder | 72 | 120 |
| 15 | - | necrosis | 0 | 120 |
| 16 | - | miscellaneous | 0 | 120 |
| 17 | 4 | no eye | 0 | 120 |
| 18 | 4 | abnormal eye size | 0 | 120 |
| 19 | 5 | abnormal tail | 0 | 120 |
| 20 | 5 | abnormal tail tip | 0 | 120 |
| 21 | 5 | abnormal tail fin | 0 | 120 |
| 22 | 5 | abnormal tail length | 0 | 120 |
| 23 | 5 | lack of somites | 0 | 120 |
| 24 | 5 | no tail detachment | 0 | 120 |
| 25 | 6 | developmental delay | 0 | 120 |
| 26 | 6 | hours post fertilization | 0 | 120 |
| 27 | 7 | deformation head | 0 | 120 |
| 28 | 7 | smaller head | 0 | 120 |
| 29 | 7 | deformation yolk sac | 0 | 120 |
| 30 | 7 | malformation sacculi / otoliths | 0 | 120 |
| 31 | 7 | modified structure of chorda | 0 | 120 |
| 32 | 7 | scoliosis | 0 | 120 |
| 33 | 7 | abnormal heart | 48 | 120 |
| 34 | 8 | shivering | 0 | 120 |
| 35 | 8 | tremor | 0 | 120 |
| 36 | 8 | no spontaneous tail contraction | 0 | 30 |
| 37 | 8 | no movement after hatching | 72 | 120 |
| 38 | 9 | decreased heartbeat | 48 | 120 |
| 39 | 9 | increased heartbeat | 48 | 120 |
| 40 | 9 | no heartbeat | 48 | 120 |
| 41 | 9 | beats per minute | 48 | 120 |
| 42 | 10 | no blood circulation | 48 | 120 |
| 43 | 10 | blood congestion | 48 | 120 |
| 44 | 11 | yolk sac edema | 24 | 120 |
| 45 | 11 | pericardium edema | 24 | 120 |
| 46 | 12 | no pigmentation | 48 | 120 |
| 47 | 12 | low pigmentation | 48 | 120 |
| 48 | 12 | increased pigmentation | 48 | 120 |
| 49 | 13 | early hatching | 48 | 72 |
| 50 | 13 | no hatching | 72 | 120 |
| 51 | 14 | no swim bladder | 72 | 120 |
| 52 | 14 | smaller swim bladder | 72 | 120 |

### The INTOB Software

INTOB is a modular software that consists of a user interface (UI), data base and an application programming interface (API). In the UI, experiments can be set up in a repeatable format. Chemicals are selected through a mirror of the CompTox database, ensuring unique identifiers for tested compounds (Williams et al., 2017). User-defined samples or chemical mixtures can also be entered. The molecular weight is automatically integrated, so that mass in grams can be converted to amount of substance in mole, and vice versa. Concentrations are defined and assigned to containers, which can be selected from predefined types (vial format or common well-type formats). Finally, relevant metadata is entered and stored alongside the experimental data (Supplementary Table 1). Observations can be recorded in a way that for each embryo effects can be assigned based on a set of defined effects that are documented within the database and amendable by the users. All data is stored in a relational database.

Data stored in INTOB can be explored through the UI by filtering experiments based on their metadata. Experiments can then be downloaded via a comma-separated value (csv) download functionality, where multiple csv files comprise all metadata, observations and physico-chemical properties pertaining to the selected experiments. Additionally, an application programming interface (API) access is offered via GraphQL and REST, to allow for an integration of INTOB into computational workflows.

INTOB includes a user management, where individual users can be assigned to groups and permissions define reading and writing rights for experiments created in certain groups. The INTOB administrator can assign roles and permissions as well as edit the set effects that can be applied to embryos. INTOB is commercially available under request (https://www.ufz.de/intob/index.php?en=51326).

### Fish Embryo Acute Toxicity Test

We used wild-type in-house strains of the zebrafish (*Danio rerio*). Fish were cultured at 26 ± 1 °C at a 14:10 hours light:dark cycle in a recirculating tank system and used according to German and European animal protection standards and approved by the Government of Saxony, Landesdirektion Leipzig, Germany (Aktenzeichen 75–9185.64).

To harvest the eggs, trays with artificial plants were placed in the tanks with male and female fish in the evening. Spawning took place shortly after the lights turned on in the morning. Eggs were collected and fertilized eggs were sorted and cultured from 2 hours post-fertilization (hpf) in control embryo medium until exposure at 26-28°C. Experiments were carried out based on OECD TG 236 (OECD, 2013), although there were differences in e.g. replicate number, exposure start, observation time points or recorded effects.

Chemical exposures were conducted at 26-28°C in different exposure vessels, such as e.g. glass vials (2ml), crystallization-dishes covered with watchmaker glasses or multi-well-plates (24-, 96-wells) covered with lids. Every experiment included controls (unexposed embryos) and treatments (exposed embryos), if solvents were used, a solvent control was always considered. Embryos were checked daily for effects on phenotype and survival with a stereomicroscope (Table 1) and data were entered into the INTOB database in parallel using a tablet/ web browser to access the INTOB User Interface (UI). All data (metadata and effect data) of each experiment were recorded by using the INTOB software.

### Data integration and selection for analysis

Next to the established direct input using INTOB on a tablet in the lab, ZFET data recorded at the UFZ during the last 20 years until December 2022 and documented in lab books and excel files were manually integrated into INTOB. For data analysis, all data recorded in INTOB before the 7th of March 2024 was used. Microscopic phenotypic observations on ZFE morphology, a summary of phenotypic effects, metadata, physico-chemical properties of chemicals, and the list of effects were downloaded through the csv download functionality in the UI. Information on the container layout including concentrations and coordinates in the well plates or vials for embryos was retrieved via the GraphQL API. All analyses described subsequently were conducted in R (v.4.3.0, (R Core Team, 2021)). GraphQL requests were sent to INTOB from R using ghql (v.0.1.0), the returned JSON (Javascript Object Notation) was converted into tabular format using jsonlite (v.1.8.8).

### Data processing and quality control

Initially, experiments with limited data (fewer than 15 embryos or only one tested experimental condition) were not considered for analysis. Subsequently, experiments were assessed for the quality of their control (non-chemical-treated) embryos. The probability of control embryos exhibiting no phenotypic morphological effects at the last time point of observation of each experiment was calculated across all experiments to be 93.6%. Assuming effects in unexposed embryos follow a binomial distribution with p(normal) = 0.936 and p(effect) = 1 – p(normal), we conducted one sided binomial tests on the final control observations for each experiment. For experiments with p-values < 0.05 we accepted the alternative hypothesis (p(normal) < 0.936) and did not include them in subsequent analyses. We want to point out that we consciously decided not to adjust p-values. Here, we are interested in the experiments for which we can not reject the null hypothesis (p(normal) = 0.936), with p-value adjustment there would be more experiments that fall into this group, potentially including more low-quality experiments (false negatives). Thus, without p-value adjustment, we aim to limit the type II error. To additionally limit the number of low-power experiments, experiments with fewer than 9 control embryos were excluded (Supplementary Figure 1).

### Data analysis

#### Dose response modeling

For all remaining single substance experiments with 0 or 24 hpf exposure start points and observations at 48, 72, 96 or 120 hpf, dose response modeling (DRM) was performed. For lethal DRM, only “coagulated” was considered as lethal, and all other effects as non-lethal. For sublethal DRM, all effects except for “normal” were considered as sublethal, i.e. the cumulative DRMs were fitted. For each substance, lethal and cumulative dose response curves were fit for all combinations of exposure start and observation time points using the drc package (v.3.0-1, (Ritz et al., 2015)). Two parameter log-logistic, Weibull 1 and Weibull 2 models were fitted and the best fitting model was chosen using Akaike’s Information Criterion (AIC). Further, models were only considered valid if their AIC was lower than the AIC of a linear model with slope = 0, their predicted EC50 was lower than the maximum tested concentration and the standard error of the EC50 was smaller than the EC50 itself.

#### Sensitivity ratio analysis

The sensitivity of all endpoints (sublethal + coagulation) was evaluated by modelling the EC50 values (considering all endpoints) and compared with the LC50 for each chemical. In line with the teratogenic index concept (Aalders et al., 2016; Selderslaghs et al., 2009), we calculated the sensitivity ratio (SR) as the quotient of LC50 to EC50 (Bittner et al., 2019). A value close to 1 suggests that sublethal or teratogenic effects occur at concentrations close to lethal ones.

#### Phenotypic fingerprints

To compare effect patterns between substances, compound-specific phenotypic fingerprints based on prevalences of individual effects were derived. To minimize the effect of differently chosen exposure concentration ranges, data from concentrations between the EC5 and the LC99, or in case the lethal DRM failed, data from concentrations larger than the EC5 were used. Normal embryos were excluded and for the remaining data, for each effect, the proportion of embryos exhibiting this effect within the selected concentrations was calculated. These vectors were clustered using hierarchical clustering (Ward method with Euclidean distance). Additionally, principal component analysis (PCA) was carried out to visualize the fingerprints and to identify important variables. Since a PCA creates a set of new variables (principal components) by applying linear transformations to the original variables, the contributions (loadings) of each original variable to the PCs can be used to assess importance of variables for explaining variance. Uniform Manifold Approximation and Projection (UMAP) was applied to the fingerprints to visualize clusters of chemicals with similar fingerprints. In line with the hierarchical clustering, the Euclidean distance measure was used.

#### Effect propagation

Data from 48 and 96 hpf observation were used to assess how different effects are correlated with each other. To ensure that the effects in individual embryos could be tracked over time, only experiments using well type containers (24-well, 48-well or 96-well plates) were considered. Only qualitative effect measures (yes/no) were used and uninformative (‘miscellaneous’, ‘no embryo’) effects were excluded. Pearson correlation coefficients and p-values were calculated between effects at 48 hpf and effects at 96 hpf. Correlations with p-values < 0.01 were considered significant. For visualization, this matrix was clustered using hierarchical clustering with Euclidean distance.

#### Literature review on ZFE toxicological phenotypes

To compare our systematic approach with data from literature we systematically collected and annotated studies from 2014 to 2023 that describe and assess morphological phenotypes in the ZFE after chemical exposure. Studies were searched using Pubmed and Science Direct, search terms consisted of combinations of organism specific terms (“Zebrafish larva”, “Zebrafish embryo”, “Danio rerio larvae”, “Danio rerio embryo”, “FET”) and effect specific terms (Supplementary Table 2). Studies were searched for keywords related to effects (Supplementary Table 2) and information was entered into a Zotero library. Effects were entered and annotated to match the effects used in INTOB (Table 1). Also, metadata was collected accordingly. This information included substance name, embryo age at observation (hpf) and the fish strain (Supplementary Table 3). We provide this dataset in JSON (Supplementary File 1) and csv format (Supplementary Table 4).

#### Data and code availability

The data used in this study is available at Zenodo (https://doi.org/10.5281/zenodo.11030299). This includes the data retrieved from the INTOB database, the code used for the analysis, as well as the results of the literature research as a Zotero library.

## Results

Data extracted from the INTOB database comprised a total of 638 experiments. After initial filtering (see methods for details), 609 experiments remained that contained 509 experiments that tested single substances, 65 mixture experiments, 34 experiments with environmental samples, and 10 experiments labeled “other”, which mainly include tests with commercial products. The dataset comprises experiments on 136 individual substances, on average a substance has been assessed in 2.6 experiments. The most often investigated substances by number of embryos tested are 3,4-dichloroaniline (a commonly used positive control substance), azinphos-methyl, and deltamethrin (Figure 2a). Most experiments made observations at three different time points after exposure, however, some experiments included up to 15 time points of observation. An overview over the most commonly used exposure start and observation time points and the associated numbers of experiments is shown in Figure 2b. On average, per experiment 7 individual exposure concentrations were tested using a total of 95 embryos, however, there is variation, as the number of exposure concentrations range from 2 to 20 and the number of embryos range from 21 to 480 (Figure 2c). In summary, this dataset presents a diverse set of experiments with varying sample sizes, exposure conditions, and coverage of observation time points per substance.

**Figure 2:**
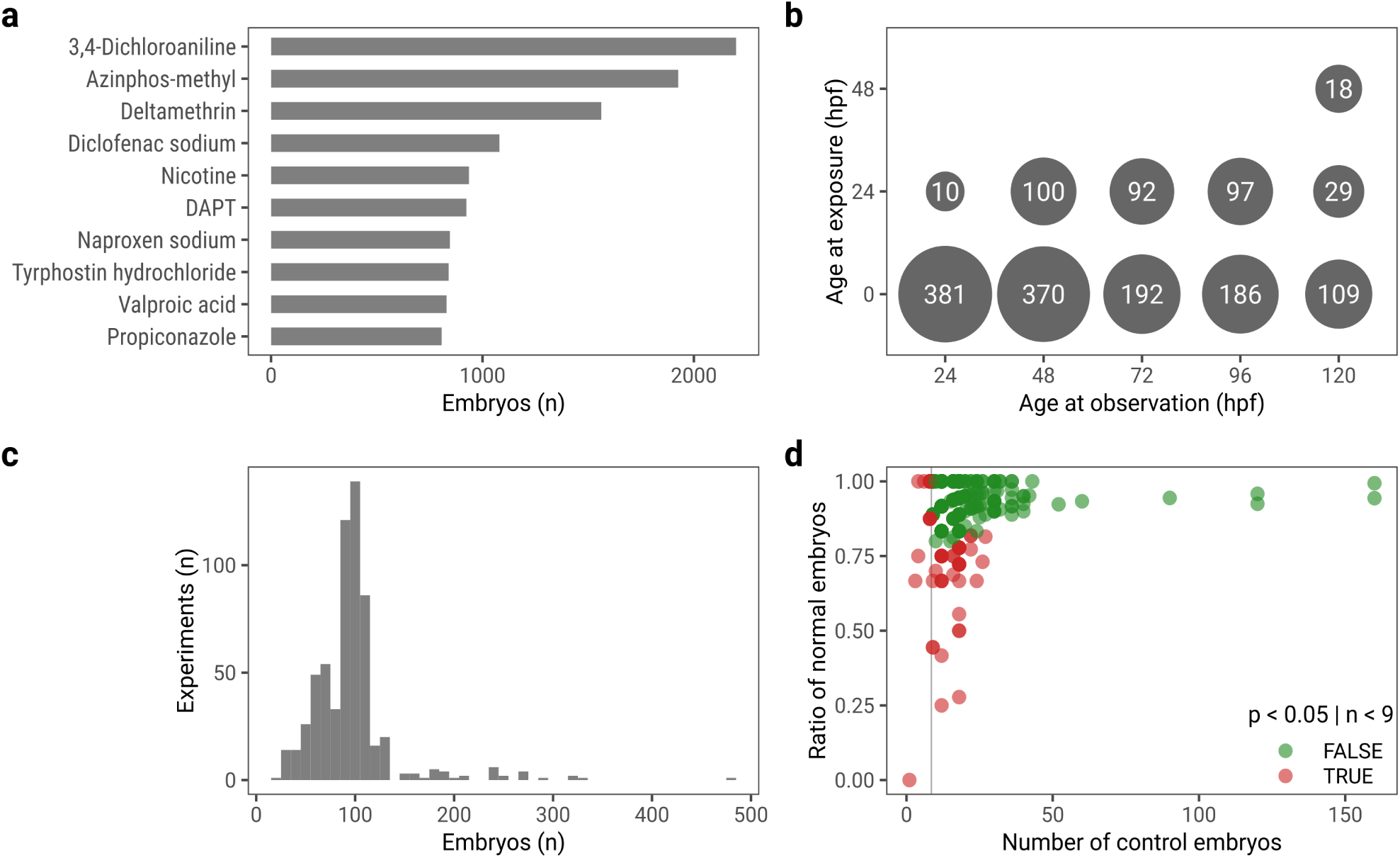
Data overview. **a**, numbers of tested embryos across all experiments for the 10 most frequently tested substances. **b**, numbers of experiments per exposure start and observation time points (most common time points depicted). **c**, histogram showing the numbers of experiments with numbers of embryos. **d**, red dots signify experiments where control embryos were identified to not stem from a binomial distribution with p(normal) = 0.936 or had fewer than 9 control embryos, green dots represent experiments for which the null hypothesis was not rejected.

### Control variability in ZFETs

Control embryos, meaning embryos that were not exposed to toxic chemicals, but exposed to media or solvent vehicles (also called negative controls or solvent controls) can exhibit morphological effects, similar to embryos exposed to chemicals. These result from biological variation and from environmental factors that are difficult to control. To account for this, experiments are typically discarded if control embryo quality does not meet a predefined cutoff. For example, the OECD test guideline 236 requires that 90% or more of the control embryos survive until the end of the experiment at 96 hpf (OECD, 2013). For experiments with defined numbers of embryos this is adequate, however with varying sample sizes this number might need statistical adjustments. For this reason, we implemented a strategy that utilizes binomial tests to determine whether the distribution of normal and non-normal control embryos in a given experiment could plausibly stem from a binomial distribution for which the probability of an embryo exhibiting no effect was calculated from the 11080 control embryos available in this dataset. Across these embryos, the rate at which unexposed embryos are “normal” at the end of an experiment is 93.6%, the rate at which they survive until the end of the experiment is 95.9%. Applying these criteria to the 609 experiments, 62 were determined to have insufficient embryo quality and were removed before further analysis, leaving 547 experiments for further investigation (Figure 2d).

### Dynamic range of effect concentrations in the ZFET

For analyzing and comparing effect concentrations we selected data from all single substance experiments that followed a 0 to 96 hpf or 24 to 96 hpf exposure design, corresponding to 71 unique substances. Out of 180 experiments fitting these criteria, 166 could be used for cumulative DRM, the remaining experiments only assessed mortality and could thus only be used for lethal DRM. Given that data was available, we estimated EC50_0-96hpf_ and EC50_24-96hpf_ as well as LC50_0-96hpf_ and LC50_24-96hpf_ for all substances (Figure 3a). We also calculated the SR, within both exposure scenarios. For 58 substances, at least one model could successfully be fitted; for 55 substances, at least one cumulative model (considering all sublethal and lethal effects) could be fitted; for 43 substances, at least one sensitivity ratio could be calculated; and for 13 substances, data was available; however, none of the models succeeded in fitting (Figure 3b). For most substances, SRs were small, only six substances had SRs above 5 (benzovindiflupyr: 22.4; valproic acid: 16.3; ferbam: 9.8; diclofenac sodium: 6.2 (0-96 hpf) and 8.4 (24-96 hpf); acetaminophen: 7.2; propoxur: 6.7).

**Figure 3:**
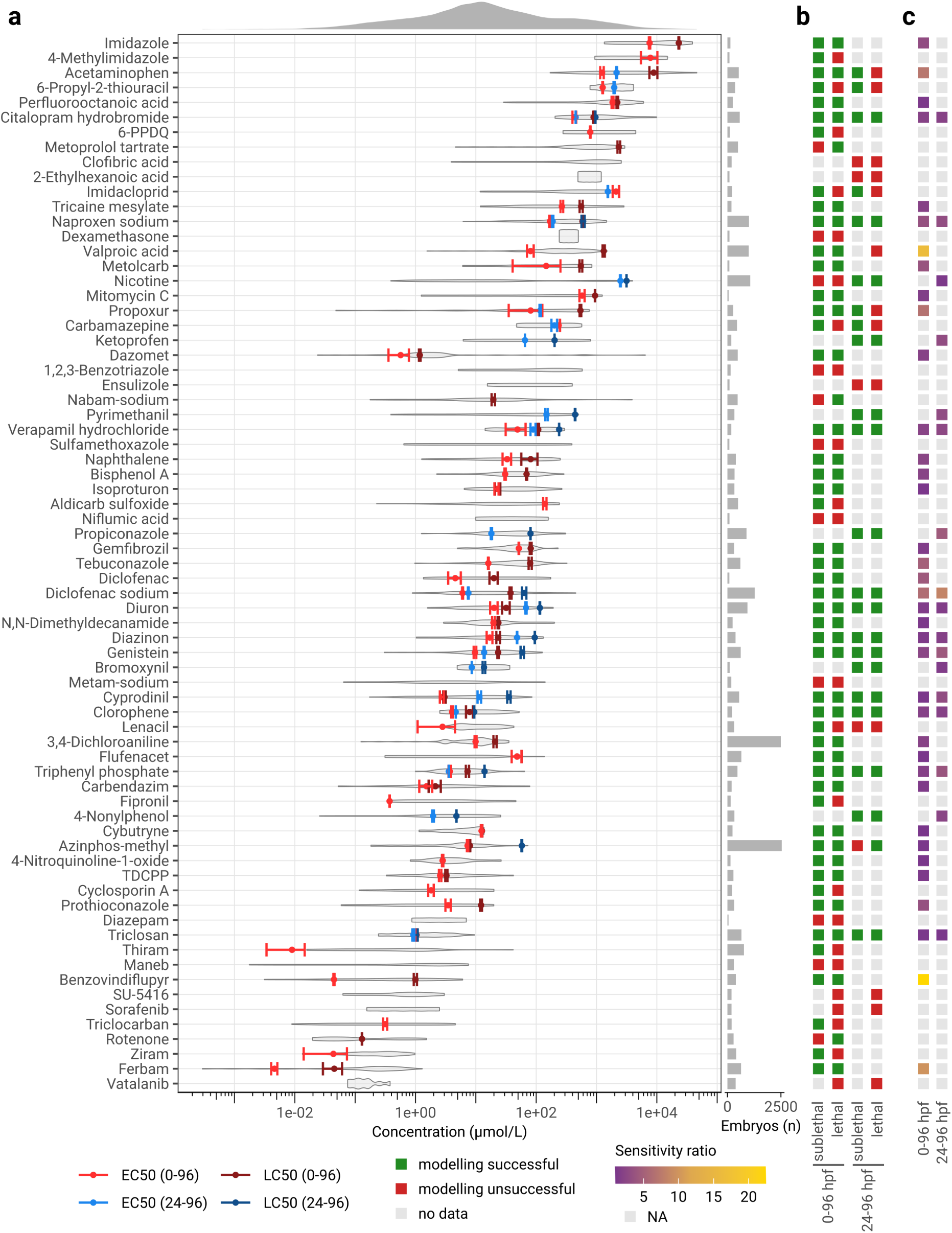
Concentration ranges for ZFET data with observations at 96 hpf. **a**, EC50 and LC50 values were modeled per compound from observational data of experiments with 0 hpf or 24 hpf exposure start points, error bars indicate their standard error. Violin plots indicate the distribution of measured concentrations. **b**, summary of data availability and modeling. **c**, sensitivity ratios (SRs). When no SR could be calculated, a gray square is displayed. Substance names from **a** carry over to **b** and **c**. Some substance names were abbreviated or changed to shorter synonyms: n-(1,3-Dimethylbutyl)-N’-phenyl-p-phenylenediamine (6-PPDQ), ethyl 3-aminobenzoate methanesulfonic acid salt (tricaine mesylate), and tris(1,3-dichloro-2-propyl) phosphate (TDCPP).

In most experiments (75%), embryos were exposed immediately after fertilization, fewer experiments initiated exposure at 24 hpf (21%). If a SR could be calculated for both exposure scenarios for one substance, these ratios were generally in agreement (Figure 3c). In a few cases, the SR was higher in experiments with a 24 hpf exposure start, notably in cyprodinil (0 hpf exposure: 1.1, 24 hpf exposure: 3.1) and genistein (0 hpf exposure: 2.4, 24 hpf exposure: 4.3).

Comparing EC50 and LC50 values across exposure start points, there are substances for which the exposure start does not influence effect concentrations, such as citalopram hydrobromide, naproxen sodium, chlorophene or triclosan. For other substances there was a time-dependent effect. For example, for verapamil hydrochloride, diclofenac sodium, diuron, diazinon, cyprodinil and genistein, toxicity was consistently higher when embryos were exposed at 0 hpf as opposed to a 24 hpf exposure start. Two examples were found (carbamazepine, imidacloprid) where later exposures (at 24 hpf) led to higher toxicity compared to early exposures (0hpf). Similarly, for nicotine concentration-dependent effects could be observed when ZFE were exposed at 24 hpf, in experiments with an exposure start at 0 hpf, no DRMs could be modeled due to low toxicity across all tested concentrations.

The concentrations across these substances depicted in Figure 3a are normal-distributed on a logarithmic scale and span 9 orders of magnitude with a range from 2.9e-04 µmol/L (1.2e-04 mg/L) to 4.6e+04 µmol/L (7.0e+03 mg/L), a mean of 3.7e+02 µmol/L (7.7e+01 mg/L) and a standard deviation of 1.9e+03 µmol/L (3.1e+02 mg/L).

### Clusters of chemicals with similar morphological effect fingerprints

For recording the impact of chemicals on the development of ZFE we defined 52 endpoints, which are listed and annotated within the INTOB database (Table 1). With this unified vocabulary, experiments are comparable with each other and effect fingerprints for individual substances can be analyzed. Therefore, we created toxicological effect fingerprints for experimental data of single substances that a) followed an exposure design of 0-96 hpf or 24-96 hpf, b) where at least a cumulative DRM could successfully be modeled, and c) that were obtained with concentrations between the EC5 and the LC99 (or above the EC5, if no lethal DRM could be modeled). For each effect, the proportion of embryos exhibiting the effect in the respective concentration range was calculated, the vector of all effect proportions defines the fingerprint for a substance. After filtering out substances where fewer than 30 embryos were available to calculate the fingerprint, as well as effects that occurred on average in less than 1% of embryos, fingerprints could be derived for 47 substances, corresponding to 133 experiments and 6225 embryos (Figure 4).

**Figure 4:**
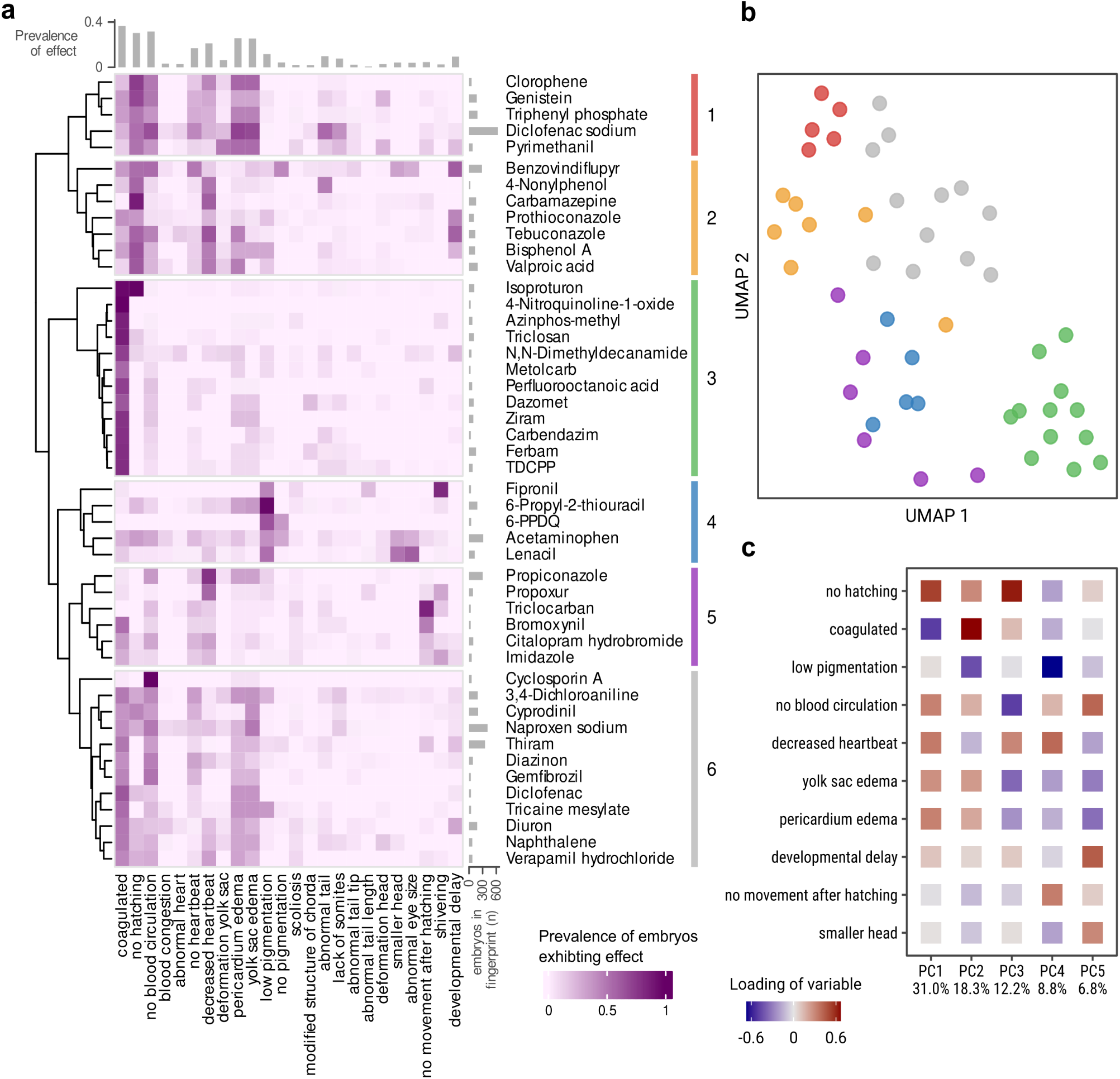
Phenotypic fingerprinting of chemical effects at 96 hpf. **a**, clustered heatmap of phenotypic fingerprints with values representing the proportion of embryos exhibiting an effect when exposed to concentrations between the EC5 and LC99 of the respective substance. Some substance names were shortened for brevity: n-(1,3-Dimethylbutyl)-N’-phenyl-p-phenylenediamine (6-PPDQ), ethyl 3-aminobenzoate methanesulfonic acid salt (tricaine mesylate), and tris(1,3-dichloro-2-propyl) phosphate (TDCPP). **b**, UMAP embedding of the fingerprints for visualization of relations between clusters. **c**, the 10 most important effects for explaining variance based on the Euclidean norm of their contributions to the first five principal components.

Clustering of the substances based on their fingerprints into six clusters revealed distinct groups ranging from 5 to 12 substances in size (Figure 4a, b). Cluster 3 (including isoproturon, ferbam and carbendazim, among others) is the most distinct cluster, defined by a coagulation phenotype. Substances in this group exhibit almost no sublethal effects. Evidently, these substances have sensitivity ratios close to 1 (Figure 3c).

„Coagulation“ is an important endpoint for explaining variance in a PCA of the fingerprints, only surpassed by „no hatching“ (Figure 4c, Supplementary Figure 5c). Interestingly, there are clusters in which substances exhibit „coagulation“ but fewer „no hatching“ effects (clusters 3 and 6, which include, among others, 3,4-dichloroaniline, naproxen, and diuron), and clusters where „no hatching“ is a predominant effect, while „coagulation“ is less prevalent (cluster 2, including, among others, bisphenol A and valproic acid, and cluster 1, which includes genistein and diclofenac sodium among other compounds). The third most important effect is „low pigmentation“, which very distinctly defines cluster 4 (including 6-propyl-2-thiouracil (6-PTU), 6-PPDQ, lenacil, acetaminophen, and fipronil), in which few other effects were observed. The fourth most important effect for explaining variance is „developmental delay“, which, however, does not seem to be important for defining the clusters. Cluster 5 (including propiconazole, imidazole, bromoxynil, among others) represents another group of substances that is defined by a small set of effects, namely the behavioral effects, no movement after hatching“ and „shivering“.

Next to effects that seem to be drivers for particular clusters, it is obvious in Figure 4a that substances in clusters 1, 2, and 6 exhibit a wide range of sublethal effects. Substances of cluster 2 cause „no hatching“, „decreased heartbeat“ and, to a lesser extent „developmental delay“ and edema effects in ZFE. Edema effects („yolk sac edema“ and „pericardium edema“) which are highly correlated, as can be seen by the direction and magnitude of their loadings in the PCA (Figure 4c, Supplementary Figure 5c), occur often after exposure with substances in clusters 1 and 6, and to some extent in cluster 2. Fingerprints for other time points were also calculated and clustered (Supplementary Figures 2-5). Comparing them, it stands out that there is always a cluster of substances defined by coagulation. It should be pointed out that individual substances often exhibit different fingerprints at different time points, indicating an effect of exposure duration.

### Occurrence and co-occurrence of morphological effects dependent on exposure time, concentration, and substance

Morphological effects observed at one time point are not independent of those of another time point in the same experiment. To determine the correlation, we calculated Pearson correlation coefficients (PCCs) between vectors of effects from all non-control embryos with observations at 48 hpf and at 96 hpf and from experiments that used well-type containers (24-well, 48-well or 96-well, equating to a total of 778 embryos, (Figure 5a). Effects at 48 hpf are typically strongly correlated with themselves at 96 hpf. One exception is „decreased heartbeat“, which is only weakly correlated (PCC = 0.14). Instead this effect, observed at 48 hpf, correlates with more severe heart-related effects at 96 hpf („no blood circulation“ and „no heartbeat“, among other effects). Other examples are “no tail detachment” and “no pigmentation”, which are not significantly correlated with themselves because they are not yet well pronounced after 48 h.

**Figure 5:**
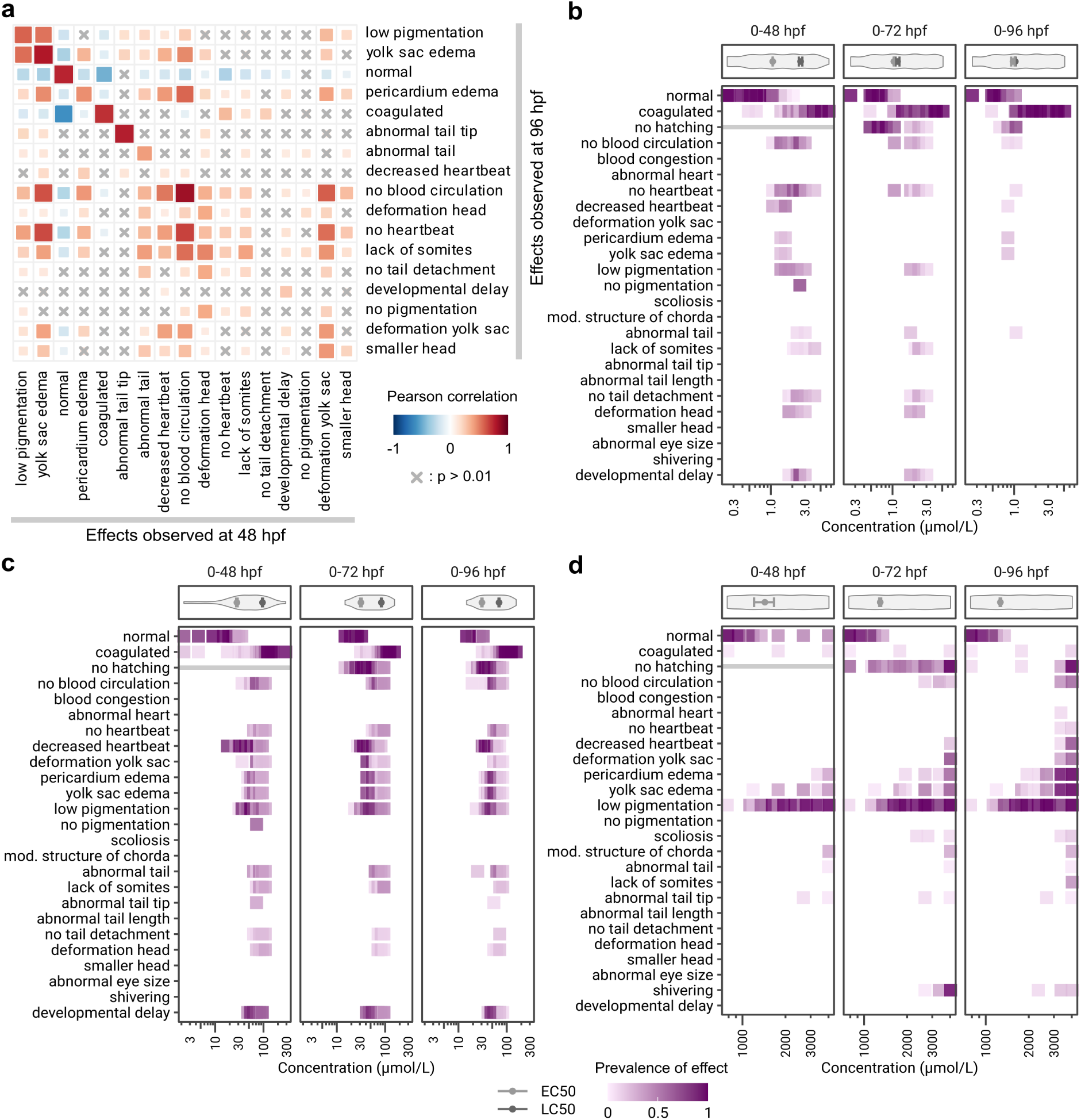
Dependency of observed effects on time and concentration. **a**, correlations between effects observed at 48 hpf and effects observed at 96 hpf in the same embryos. Correlated effects are marked in red, anti-correlated effects in blue, insignificant (p > 0.01) correlations are marked with an x. **b**, **c**, **d**, detailed phenotypic fingerprints of **b**) triclosan, **c**) bisphenol A, and **d**) 6-propyl-2-thiouracil (6-PTU), respectively. Panels on top of the fingerprints display the distribution of applied concentrations and modeled EC50 and LC50 values, error bars indicate the respective standard error. Effects that are not valid at a certain time point are marked with a gray line.

PCCs between effects at 48 hpf and the „coagulation“ effect at 96 hpf are almost all not significant (p > 0.01), exceptions are a strong and logical anti-correlation with „normal“ and weak but significant correlations to the three effects specified as lethal in the OECD TG 236, namely „no heartbeat“, „lack of somites“, and „no tail detachment“.

Among all chemical-exposed ZFE (n = 90972) that were considered in this analysis, 53.4% were found to be not affected by the treatment and tagged as “normal”. The term „normal“ is anti-correlated with all effects except itself, indicating that a large proportion of unaffected embryos do not exhibit any adverse effects over time. Additionally, „normal“ at 48 hpf and „coagulated“ at 96 hpf have the strongest negative correlation of all combinations of effects (PCC = -0.61), indicating that „normal“ embryos rarely transition to „coagulated“. Other prominent correlations were observed in cases where 48 hpf embryos were classified as developmentally delayed. Effects at 96 hpf strongly correlated to this were, apart from „developmental delay“ itself, „no pigmentation“, „no tail detachment“, „deformation head“, and „no blood circulation“.

Detailed phenotypic fingerprints reveal that individual effects occur in a time and concentration dependent manner. Figure 5b, c, d show three examples with different effect patterns over time and concentration ranges. Observations at 48, 72 and 96 hpf following exposure to triclosan (Figure 5b) exhibit a pronounced effect of exposure duration on the sensitivity ratio. At 48 hpf a range of sublethal effects could be observed, determining the EC50 at that time point, before coagulation set in at higher concentrations, determining the LC50. At 96 hpf this changes and the distribution of effects is almost binary, where embryos exposed to low concentrations were normal and embryos exposed to higher concentrations coagulated, resulting in a sensitivity ratio of close to 1 (Figure 5b). In contrast, effect patterns following bisphenol A exposure did not change over time and were comparatively constant (Figure 5c). A concentration dependency can be observed for sublethal effects, where the prevalence of an effect increases with concentration. With higher concentrations the prevalence of coagulation increases with the result that the sublethal effects decrease. It is also obvious that some sublethal effects are more sensitive than others, for example “decreased heartbeat” could be observed at lower concentrations compared to the more severe effects like “no heartbeat” or abnormal tail effects (Figure 5c). In exposures with 6-PTU (Figure 5d), concentrations were not high enough to reach lethality and the observed effect patterns are defined by the same set of sublethal effects across all time points. The most frequent effect was “low pigmentation”, which could be observed at concentrations well below concentrations where other sublethal effects started to emerge. For the two edema effects (“pericardium edema” and “yolk sac edema”), additionally to a concentration dependency, an increase of these effects with time could be observed. The EC50 value stays almost constant over time as only the number of embryos exhibiting any sublethal effect are counted. This shows how standardized data and a comprehensive and comparable analysis enlarges the amount of interpretable information.

### Comparison of chemical effects in ZFET based on data from literature

In a final step we aimed to compare our data with data from literature. Therefore, we conducted an extensive literature review of studies assessing ZFE phenotypic effects within a toxicological context published between 2014 and 2023. We obtained 162 individual publications and a total of 522 distinct chemicals, samples or mixtures that were investigated in those studies. The heterogeneity across all the studies is extremely large with variation in the considered number of effects, time points and exposure windows, and the selection of investigated concentrations (Supplementary File 1, Supplementary Table 4).

We selected substances a) for which observations at 72, 96 and/or 120 hpf were available and b) that were described with observations in at least two publications. In most cases observations are reported in a “yes/no” manner, not allowing for quantitative prioritization of effects. We transferred descriptions on effects to a standardized vocabulary and performed clustering of compounds according to their phenotypic fingerprints that could be generated from the studies. Hierarchical clustering revealed that substances tested in separate studies rarely clustered together, clusters were rather defined by individual studies (Supplementary Figure 6a).

Based on the selected criteria the overlap of substances investigated in our study and those where data from literature was available was small. One example, where comparison to our data was possible is valproic acid. Brotzmann and colleagues performed two studies investigating the effects of valproic acid and different analogues, one study in 2021 with 9 analogues, and another study in 2022 with 14 analogues (Brotzmann et al., 2022, 2021). Although substances from these studies presumably act in similar ways, their effect patterns are hardly comparable due to different methodological approaches within the two different studies. Comparing the fingerprints of valproic acid with our data shows some overlaps but no clear picture (Supplementary Figure 6b). Reported EC50 and LC50 values differ across studies, but whether this is due to data analysis, modeling strategy, or experimental differences is difficult to evaluate (Supplementary Figure 6c).

## Discussion

In the present study, we analyzed a dataset of ZFET data containing over 600 experiments. We provide a method to assess control variability in ZFETs taking into account varying sample sizes. We show that morphological effect fingerprints can be used to effectively cluster substances, and that factors such as exposure start or observation time points can have an influence on such fingerprints. These types of analyses were made possible with the development of INTOB, a tool that stores ZFET data in a machine-readable, consistent format. For further development and use, we provide both, the ZFET data with observations for 136 chemicals as well as the data analysis workflows via Zenodo (https://doi.org/10.5281/zenodo.11030299).

Morphological effect fingerprints might be used in the future to infer mechanisms of toxic action, if they can be linked to specific phenotypic patterns. Our study provides evidence that hierarchical clustering effectively groups substances with similar phenotypic fingerprints, highlighting relationships between chemical exposure and observed developmental effects in embryos. Truong and colleagues already demonstrated that simultaneous evaluation of different phenotypic and behavioral endpoints of the zebrafish embryo revealed distinct patterns of chemical responses, aiding in the identification of mechanistic pathways (Truong et al., 2014). In our study, we found a cluster of substances defined by a low pigmentation phenotype. This cluster includes 6-PTU, a model compound for developmental toxicity and inhibitor of the thyroid peroxidase, which is involved in the synthesis of the thyroid hormones (TH) triiodothyronine (T3) and thyroxine (T4). A link between thyroid hormone levels and pigmentation has been observed and a mechanism has been suggested where TH regulates the differentiation of the different pigment cells in zebrafish (Saunders et al., 2019). Walpita and colleagues observed low pigmentation in ZFE, when levels of T3 where lowered by antisense oligonucleotide targeting of the type 2 iodothyronine deiodinase, which catalyzes the conversion of T4 to T3, a phenotype they were able to rescue by exogenous exposure with T3 (Walpita et al., 2009). Indeed, other substances of this cluster (see Figure 4) have also been indicated as disruptors of the thyroid hormone system. 6-PPD and its metabolite 6-PPDQ and their effects on the thyroid hormone system have been described and mechanisms of action were hypothesized (Bohara et al., 2024; Peng et al., 2022; Zhang et al., 2023). For acetaminophen low pigmentation phenotypes have been observed and transcriptomic analyses indicate involvement of thyroidal pathways (David and Pancharatna, 2009; Wang et al., 2024). Exposure to fipronil decreased T3 and T4 levels in adult zebrafish and their offspring (Xu et al., 2019), presumably by affecting key genes involved in TH synthesis and regulation (Ma et al., 2024). Conversely, for lenacil the European Food Safety Authority (EFSA) concluded that endocrine disruption criteria are not met in the T-modality, however no results from ZFE were discussed in this review (Álvarez et al., 2024).

The disruption of the thyroid system during zebrafish development has not only been shown to reduce pigmentation but also the size of the eyes (Baumann et al., 2016). For acetaminophen and lenacil, we could observe effects on head and eye size, albeit not for the other substances in this cluster (Fipronil, 6-PTU, 6-PPDQ). Benzovindiflupyr exposures also caused decreased head and eye sizes as well as low pigmentation effects, however, in the clustering it is grouped with other chemicals as it also exhibits many other effects. These findings show the limitations of the current approach, but, at the same time highlight the potential for further mechanism-related studies and generating mechanistic evidence, e.g. on modes of action by read-across. Many effects are not independent of each other, but instead often co-occur or transition into related effects across observations. Investigation of these trajectories, possibly with higher temporal resolution, could give further insight into the ways at which chemicals affect ZFE. In some cases, like triclosan, these time-dependent changes can result in drastically different phenotypic fingerprints for different observation time points (see Figure 5b). Including information on such dynamics in analyses might be crucial to detect patterns related to specific mechanisms. In this line, specificity of phenotypes might also be considered in more detail in the future as less specific effects might obscure MoA-related patterns and obstruct the clustering in the current approach.

During the course of the 96 h ZFET, the ZFE develops a liver and gastrointestinal system, which enables active metabolization and biotransformation of chemicals. This can result in a detoxification of a compound or the formation of toxic metabolites. Increasing or decreasing EC or LC values over time or within different exposure windows are indicative for such processes, which provides mechanistic evidence for risk assessments. Here, we showed that lower effect concentrations were more often found when exposures start early after fertilization compared to later exposures, possibly indicating detoxification processes in later embryonic stages due to an active metabolism. The opposite was observed for some compounds (carbamazepine, imidacloprid and nicotine) that are known act on the nervous system. The nervous system is developed rather late in ZFE, which may explain the higher toxicity in later stages (Blader and Strähle, 2000). This gap of the OECD TG 236, which requires an exposure start directly after fertilization, was also highlighted by Sobanska and colleagues when LC50 values of ZFETs were compared with those of adult fish (Sobanska et al., 2018). Therefore, we conclude that adding a second window with an exposure start at 24 hpf to the OECD TG 236 would increase the potential of the assay to provide more mechanistic evidence and and to completely avoid animal testing with adult fish in the future.

Comparing sublethal and lethal effect concentrations helps to identify chemicals that have the potential to be teratogenic and/or developmentally toxic (Aalders et al., 2016; Selderslaghs et al., 2009; Teixidó et al., 2022). In our study, high SRs were found, e.g. for propoxur and valproic acid, two chemicals where an adverse impact on the health of the fetus due to exposure during pregnancy have been described (Andrade, 2018; Ostrea et al., 2012). Another chemical with a high ratio between LC50 and EC50 is benzovindiflupyr, a rather novel fungicide for which teratogenicity has not yet been described. Using the INTOB database with additional and quantitative endpoints from e.g. automated imaging of morphology or behavior, in larger systematic studies that include well-known teratogens can provide further insight into the potential of the ZFE as a valid prescreening tool for teratogenicity.

One of the limitations of phenotypic assessments is that even users that received the same training judge observed effects differently. A harmonized vocabulary is a step forward to improve reports, and the implementation of ontologies for toxicological screens can reduce the variability in reported phenotypes and can enhance agreement and consistency across laboratories (Thessen et al., 2022). Biases induced by experimenters due to judgment of effects in microscopic observations can be reduced by incorporating data from automated imaging systems and respective software tools as have already been established for the zebrafish embryo model (Nöth et al., 2024; Teixidó et al., 2022, 2019). Such datasets provide more and quantitative features and could be extended further with readouts other than morphology, such as behavioral data. The current version of the INTOB software already enables upload of such data, providing the potential for larger and more detailed analyses in the future and enabling comparative assessments of chemicals, also in the context of high throughput testing.

Structured and machine-readable data in toxicology is key to accelerate the assessment of chemicals and apply read-across assessments. While databases, such as the US EPA ECOTOX DB or the US EPA CompTox dashboard provide data on EC and LC values, mechanistic information on morphological phenotypes are rarely reported (Olker et al., 2022; Williams et al., 2017). Within the ZFIN (https://zfin.org/, (Bradford et al., 2022)) a repository zebrafish phenotypes has been annotated with an ontology that is much broader than the one we used in this study. While phenotypes reported in literature for chemical exposed zebrafish are listed in the ZFIN repository, information on experimental metadata are not provided. Similarly, our own literature review revealed that gaining insights based on data from literature is tedious, as information is often not available in machine readable formats and effects are not described using a unified vocabulary, resulting in poor interoperability and comparability. Using the INTOB software helps to overcome these limitations and enables improved data management of toxicological data including metadata. Via a GraphQL API, the software enables a quick data export in a machine-readable format, e.g. for uploading them into larger repositories, or to run customized analyses.

Besides individual chemicals, the software enables data input for chemical mixtures and complex samples. With this, analysis and comparison of individual chemicals with mixtures is easily possible in terms of quality and quantity of effects and effect concentrations. Complex mixtures in environmental samples contain up to several hundred chemicals (Finckh et al., 2024, 2022) that occur in the aquatic environment across 6 to 7 orders of magnitude from the ng/L to the mg/L range (Busch et al., 2016; Finckh et al., 2022; Halbach et al., 2021). With our study we could show that the range of concentrations causing sublethal to lethal effects in ZFE for a diverse set of chemicals covers 7 orders of magnitude from the low µg/L to high mg/L ranges. With this, our study reveals for the first time, the suitability of the test system ZFE to cover a wide range of concentrations that are relevant in terms of real-life exposures.

In this study we aimed to investigate the potential of phenotypic fingerprints based on structured and machine-readable ZFET data and metadata, and decided on a particular way how to integrate the data for the analyses. We developed fingerprints for specific time points and integrated observations across different concentrations. The database is structured in a way that multi-dimensional analyses considering at least concentrations and time for each observation and effect endpoint is possible once the experiments are set up in ways to systematically gain this data. Such data-cube-based analysis can be established in the future to inform and improve chemical risk assessment. Furthermore, toxicological data provided in a FAIR way can be integrated into larger assessment schemes and data networks applied to train AI models on our way towards an *in silico* toxicology.

## Supporting information

SI_file1_zotero_phenotype_library_json

SI_tab4_zotero_phenotype_library_csv

## Acknowledgments

The development of the INTOB software was supported and funded by the DataHub initiative of the Research Field Earth and Environment of the Helmholtz Association and the Federal Ministry of Education and Research (BMBF), by the UFZ via the INTOB “transmarket” project, a technology transfer project within the “transfun” program, and by the BMBF project DiMEP (FKZ: 16LW0475K). RM was funded by the NFDI4Bioimage project (Project number 501864659) funded by the German Research Foundation (DFG). Data was provided and generated within different periods of the program-oriented funding (POF) of the Helmholtz Research Program Earth and Environment (POF II to POF IV (Topic 9)). We thank Ulrike Schlägel for support in statistical questions, Paula Klein for data import, and Rebekka Lange for support in preparation of figures. We acknowledge our subcontractor, the InfAI GmbH Leipzig, for software development and programming.

## Supplementary files

- Raw data and code at Zenodo (https://doi.org/10.5281/zenodo.11030299)
- supplementary_information.pdf
- SI_file1_zotero_phenotype_library.json
- SI_tab4_zotero_phenotype_library.csv

