## supplementary_information for "Leveraging zebrafish embryo phenotypic observations to advance data-driven analyses in toxicology"

Supplementary Table 1: Experimental metadata information in INTOB.

| **Metadata information** | **Variable #1** | **Variable #2** | **Variable #3** | **Variable #4** |
|---|---|---|---|---|
| Substance | Name | DTXSID | Molecular Mass | - |
| Test concentrations | Number of concentrations | Test concentrations | Unit | - |
| Fish strain | Name of fish strain | - | - | - |
| Exposure volume | Volume per embryo in ml | - | - | - |
| Exposure temperature | Temperature in °C | - | - | - |
| Medium | Name | - | - | - |
| pH in medium at start | pH Value | - | - | - |
| Buffer | Name of buffer | Concentration in mmol/L | - | - |
| Solvent | Yes/No | Name | DTXSID | proportion in test % (v/v) |
| Experimenter | Name of experimenter | - | - | - |
| Exposure start | Age at exposure start in hpf | - | - | - |
| Additional information | Free text information | - | - | - |
| Plate configuration | Type (well plate format; Exposure vial) | Number of well plates/Vials | If vial, number of embryos per vial | - |

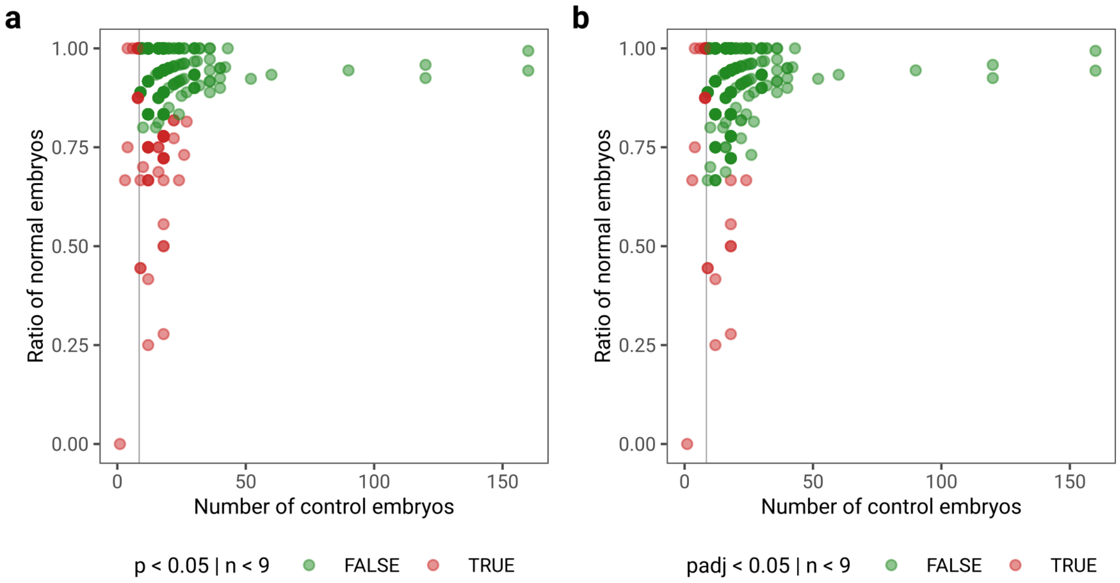


Supplementary Figure 1: Assessment of embryo quality with and without p-value adjustment. Green and red dots represent experiments that are identified as high- or low-quality experiments, respectively. The vertical line represents a cutoff for low sample size at 9 embryos. **a**, experiments with p-values < 0.05 in binomial tests or fewer than 9 embryos were identified as low-quality experiments. **b**, with p-value adjustment (Benjamini & Hochberg), the tests are more stringent in identifying low-quality experiments. In turn, they are potentially more lenient towards false negatives, i.e. low-quality experiments that are not identified as such.

Supplementary Table 2: Effect vocabulary used in the literature review.

| Observation | Abbreviation | Keywords |
|---|---|---|
| Abnormal Behavior | AB | behavior, (tail) contraction, movement, shivering, tremor, immobility, shoaling, scototaxis, thigmotaxis, sensitivity, swimming, activity, photomotor, locomotor, response, velocity |
| Abnormal Blood Circulation | AC | blood, circulation, congestion, flow, pooling, hemoglobin |
| Abnormal Eye | AE | eye, ocular, retina, optic |
| Abnormal Hatching | AH | hatching |
| Abnormal Intestine | AI | intestine, gut |
| Abnormal Pigmentation | AP | pigmentation, chromatophore |
| Abnormal Swim Bladder | AS | swim bladder, inflation |
| Abnormal Tail Effects | AT | tail, detachment, somites, fin, dorsal, caudal, pectoral, pelvic, anal, tip |
| Coagulation | CO | coagulation, apoptosis, necrosis |
| Developmental Delay | DD | development, retardation |
| Edema | ED | edema, pericardial, yolk (sac) |
| Gene Regulation | GR | gene, expression, regulation |
| Heart | HE | heart, beat, cardiac |
| Malformation | MA | head, scoliosis, lordosis, kyphosis, chorda, sacculi/otoliths, octavolateral, snout, body, axis, jaw, spine, arches, brain, cartilage, skeletal, liver, hepar, haemorrhage, facial, trabecular, palate, epiboly, trunk, pancreas, endoderm, kidney |
| Mortality Rate | MR | mortality, survival, LC, NOEC, LOEC, NOAEL, LEL, LEC |
| Neuronal Defects | ND | neurons, neurotransmitter, memory, cognitive |

Supplementary Table 3: Columns used in the Zotero library to store information.

| Zotero column name | Value |
|---|---|
| Call Number | Name of substance tested (IUPAC or common abbreviation), substances in mixtures are separated with semicolons |
| Archive | Observed effects, comma separated |
| Location in Archive | Time points at which observations were made in hours post fertilization (hpf), comma separated |
| Series Title | Fish strain, assumed WT if not otherwise specified |

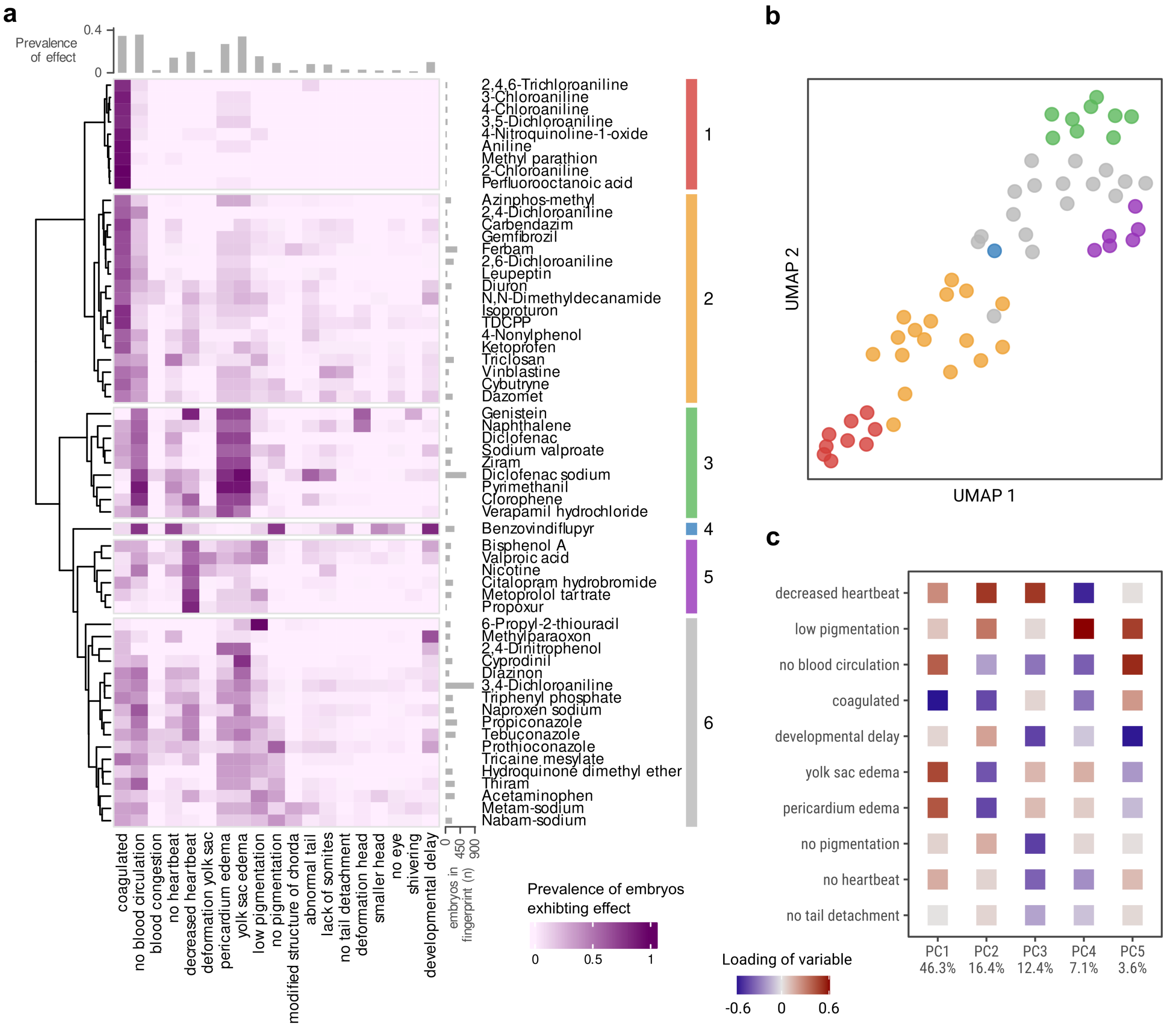


Supplementary Figure 2: Phenotypic fingerprinting of chemical effects at 48 hpf. **a**, clustered heatmap of phenotypic fingerprints with values representing the proportion of embryos exhibiting an effect when exposed to concentrations between the EC5 and LC99 of the respective substance. Some substance names were shortened for brevity: n-(1,3-Dimethylbutyl)-N'-phenyl-p-phenylenediamine (6-PPDQ), ethyl 3-aminobenzoate methanesulfonic acid salt (tricaine mesylate), and tris(1,3-dichloro-2-propyl) phosphate (TDCPP). **b**, UMAP embedding of the fingerprints for visualisation of relations between clusters. **c**, the 10 most important effects for explaining variance based on the Euclidean norm of the first five principal components.

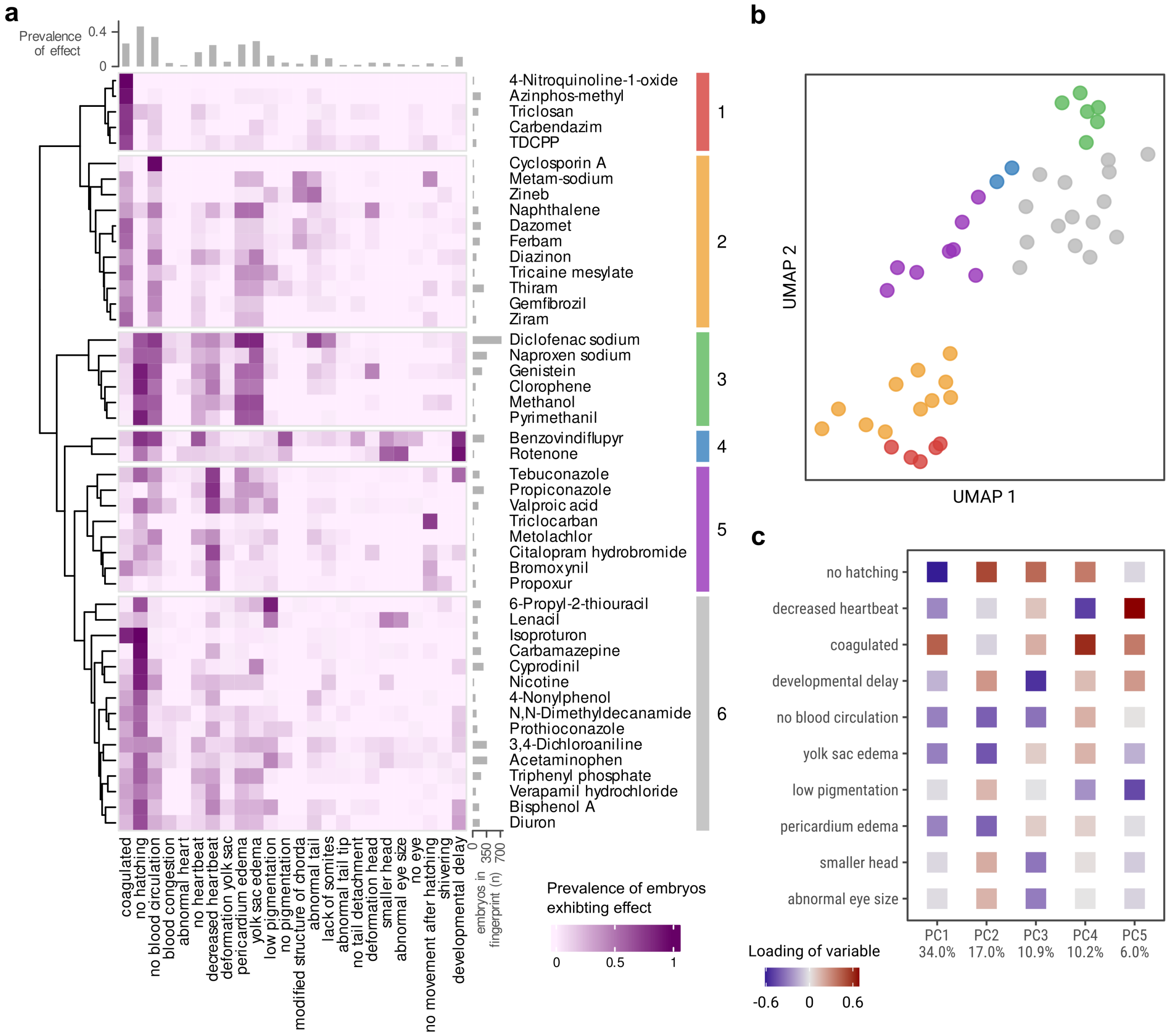


Supplementary Figure 3: Phenotypic fingerprinting of chemical effects at 72 hpf. **a**, clustered heatmap of phenotypic fingerprints with values representing the proportion of embryos exhibiting an effect when exposed to concentrations between the EC5 and LC99 of the respective substance. Some substance names were shortened for brevity: n-(1,3-Dimethylbutyl)-N'-phenyl-p-phenylenediamine (6-PPDQ), ethyl 3-aminobenzoate methanesulfonic acid salt (tricaine mesylate), and tris(1,3-dichloro-2-propyl) phosphate (TDCPP). **b**, UMAP embedding of the fingerprints for visualisation of relations between clusters. **c**, the 10 most important effects for explaining variance based on the Euclidean norm of the first five principal components.

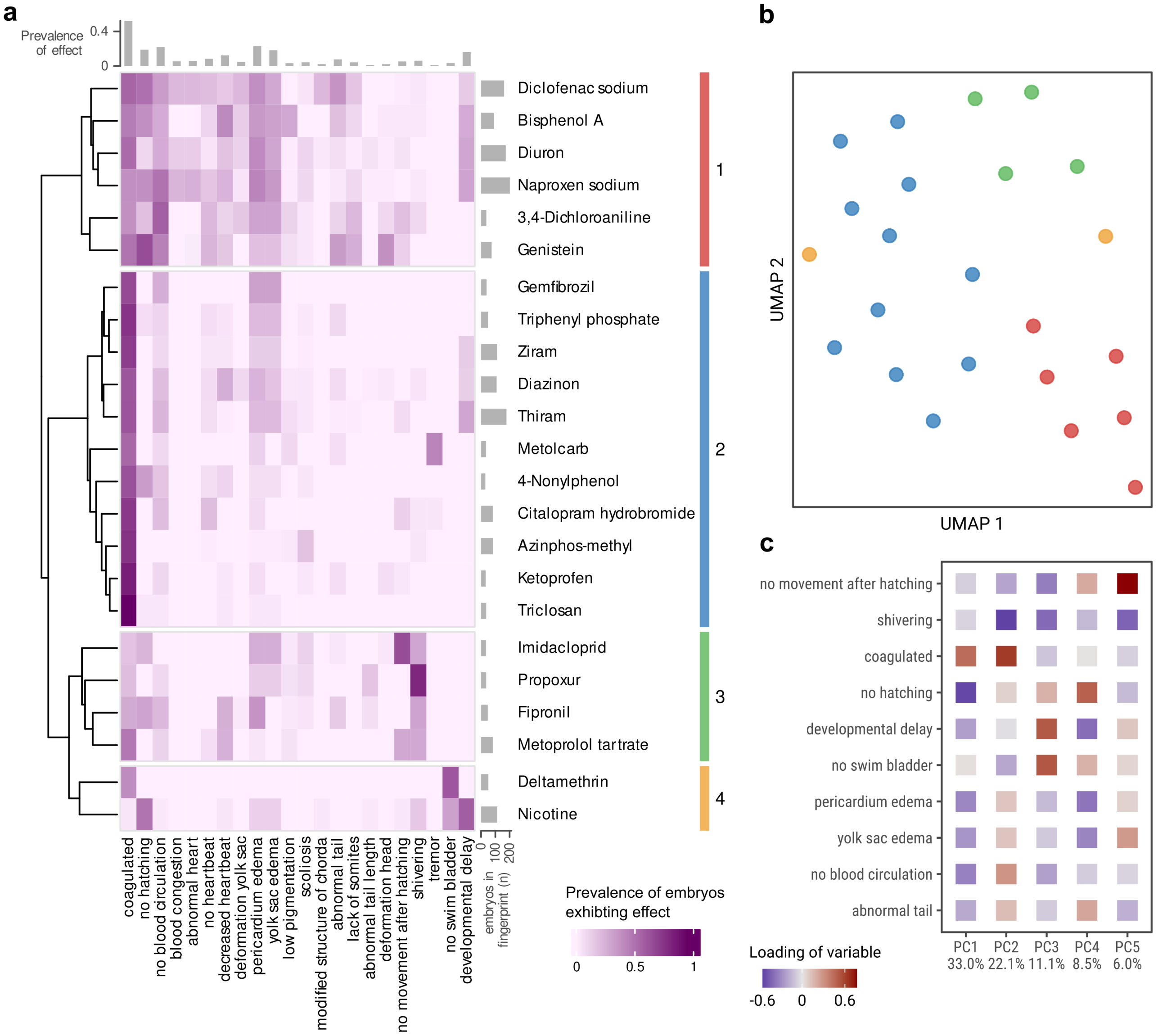


Supplementary Figure 4: Phenotypic fingerprinting of chemical effects at 120 hpf. **a**, clustered heatmap of phenotypic fingerprints with values representing the proportion of embryos exhibiting an effect when exposed to concentrations between the EC5 and LC99 of the respective substance. Some substance names were shortened for brevity: n-(1,3-Dimethylbutyl)-N'-phenyl-p-phenylenediamine (6-PPDQ), ethyl 3-aminobenzoate methanesulfonic acid salt (tricaine mesylate), and tris(1,3-dichloro-2-propyl) phosphate (TDCPP). **b**, UMAP embedding of the fingerprints for visualisation of relations between clusters. **c**, the 10 most important effects for explaining variance based on the Euclidean norm of the first five principal components.

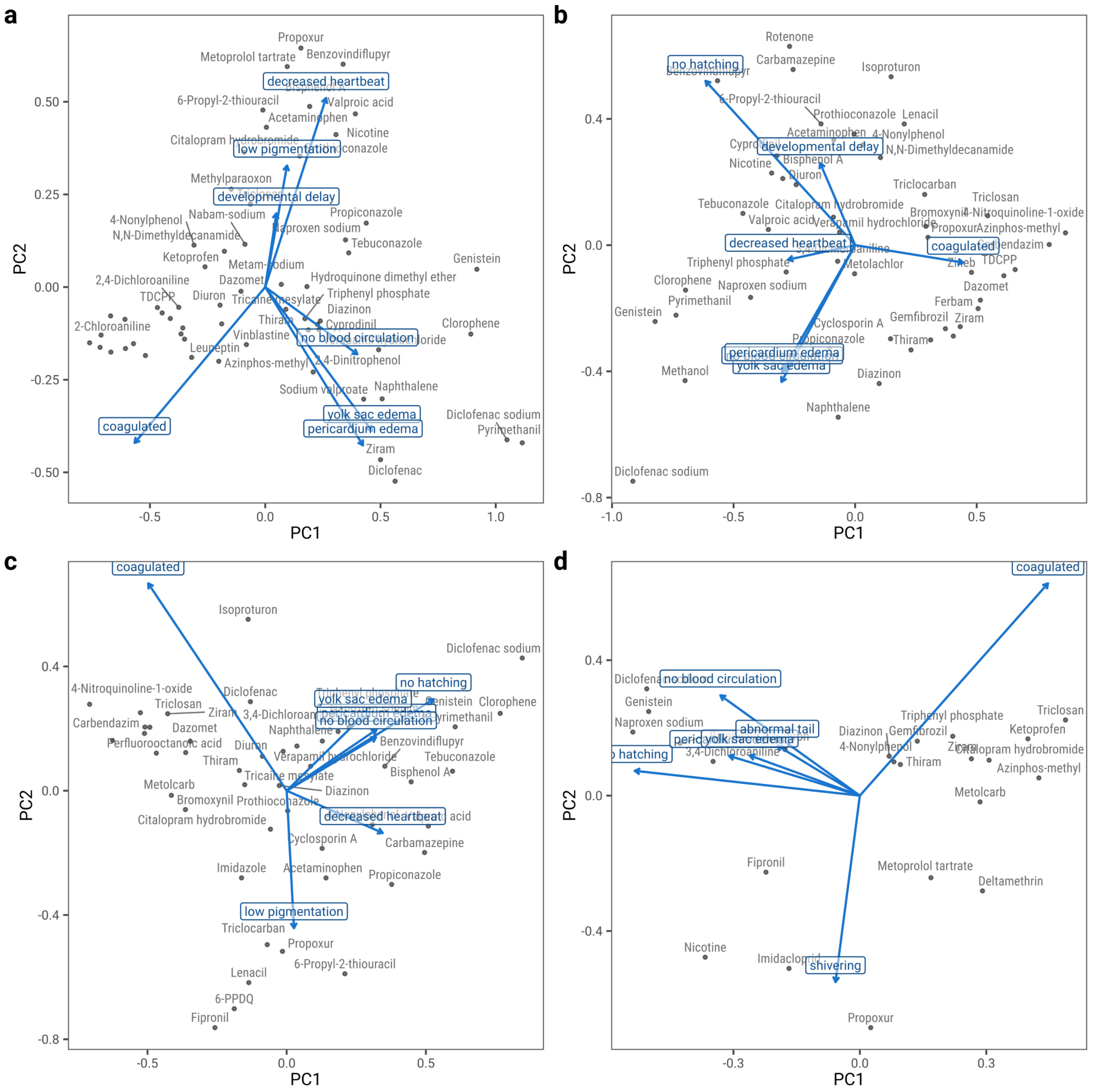


Supplementary Figure 5: PCAs of phenotypic fingerprints. Magnitude and direction of the loadings in the first two principal components for the 7 most important variables are shown in blue. Some points are not labeled due to too many overlaps. Some substance names were abbreviated or changed to shorter synonyms: N-(1,3-Dimethylbutyl)-N'-phenyl-p-phenylenediamine (6-PPDQ), ethyl 3-aminobenzoate methanesulfonic acid salt (tricaine mesylate) and tris(1,3-dichloro-2-propyl) phosphate (TDCPP). **a**, **b**, **c** and **d** show PCAs for 48, 72, 96 and 120 hpf, respectively.

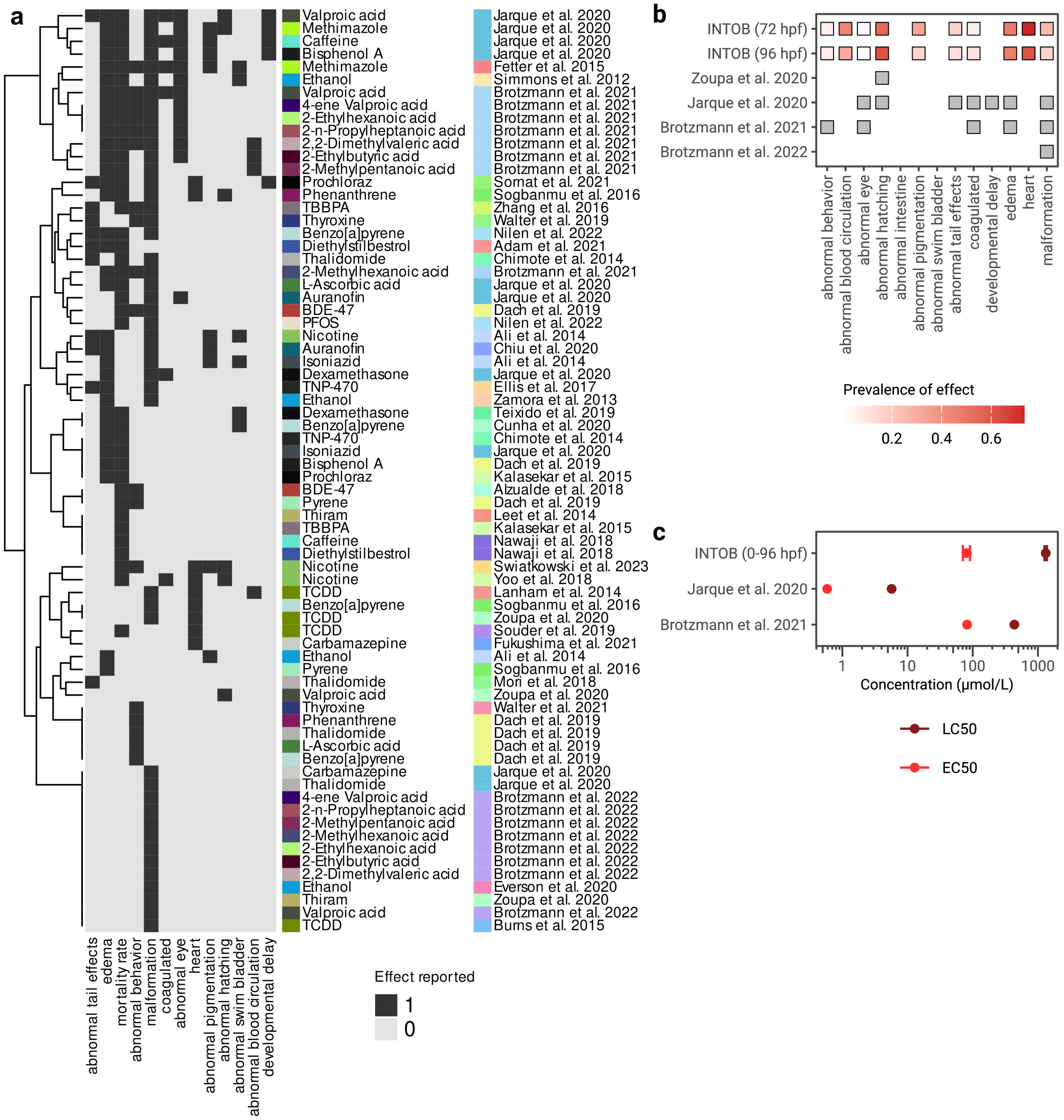


Supplementary Figure 6: Comparison of effect patterns from literature. **a**, substances assessed in more than one study and with observations at 72, 96 and/or 120 hpf are compared and clustered by hierarchical clustering. Abbreviations: tetrabromobisphenol A (TBBPA), 2,2',4,4'-tetrabromodiphenyl ether (BDE-47), O-(chloroacetylcarbamoyl)fumagillol (TNP-470), perfluorooctanesulfonic acid (PFOS), 2,3,7,8-tetrachlorodibenzo-P-dioxin (TCDD). **b**, phenotypic fingerprints from INTOB observational data and effect patterns from literature for valproic acid. **c**, EC50 and LC50 values for valproic acid modeled from INTOB observational data and reported in literature.
